# Quantitative mapping of nanoscale EGFR–Grb2 assemblies by DNA-PAINT

**DOI:** 10.64898/2026.02.16.706070

**Authors:** Alexandra Kaminer, Yunqing Li, Hans-Dieter Barth, Marina S. Dietz, Mike Heilemann

## Abstract

Receptor tyrosine kinase signaling is initiated by extracellular ligand binding, which drives the formation of membrane-protein assemblies that activate intracellular signal transduction. Accurately resolving the molecular composition of these assemblies *in situ* remains challenging due to their nanoscale dimensions and intrinsic heterogeneity. Here, we introduce a single-molecule super-resolution imaging and analysis workflow designed to resolve and quantitatively characterize individual membrane-protein assembly sites in cells. We apply this approach to the nanoscale organization of the epidermal growth factor receptor (EGFR) and its adaptor protein Grb2 following stimulation with the native ligand epidermal growth factor (EGF). As activation progresses, we observe a reduction in EGFR density at the plasma membrane, a progressive accumulation of Grb2 at EGFR assembly sites, and an increase in both dimeric and higher-order oligomeric EGFR. The experimental and analytical framework presented here is broadly applicable to the study of diverse membrane-protein assemblies.

## Introduction

The epidermal growth factor receptor (EGFR) belongs to the superfamily of receptor tyrosine kinases (RTKs) and is expressed in various human cell types, regulating key cellular processes including proliferation, differentiation, migration, and apoptosis.^[1,2]^ Dysregulation and malfunction of EGFR are associated with multiple types of human cancer, which makes it a major therapeutic target in cancer therapy.^[3–5]^ Upon ligand stimulation, EGFR dimerizes and undergoes autophosphorylation at tyrosine residues in its cytoplasmic C-terminal tail. These phosphorylation events recruit intracellular effector proteins and initiate various downstream signaling pathways.^[6]^ Among these, the mitogen-activated protein kinase (MAPK) signaling pathway is overactivated in almost one-third of all cancers,^[7]^ making it an important pathway to study at the molecular level in the context of EGFR activation. Growth factor receptor-bound protein 2 (Grb2) is the first effector of the MAPK signaling pathway recruited to activated EGFR. A reconstitution study has shown that the interaction and condensation between EGFR and Grb2 are essential for MAPK activation.^[8]^ However, how EGFR and Grb2 spatially organize and correlate *in situ* within cells remains insufficiently understood.

How signaling proteins organize into functional protein assemblies is ideally studied *in situ* in the native cellular context.^[9]^ To resolve the organization and heterogeneity of these nanoscale assemblies, high-resolution optical microscopy methods such as single-molecule localization microscopy (SMLM) have proven powerful.^[10,11]^ For example, studies employing quantitative super-resolution microscopy reported on the oligomerization of membrane receptors in general^[12–14]^ and of RTKs such as EGFR,^[15–17]^ MET,^[18,19]^ and fibroblast growth factor receptors.^[20]^ So far, these studies have examined the assembly of relevant receptors under varying ligand stimulation conditions. Beyond receptor-ligand assemblies, the formation of “signaling hubs” or protein condensates also comprises downstream effector molecules for initiating downstream signaling pathways,^[21]^ which so far remain understudied.

Here, we report the nanoscale organization of EGFR and its adaptor protein Grb2 following EGF stimulation over time *in situ*. We recorded two-target SMLM data using DNA-points accumulation for imaging in nanoscale topography (DNA-PAINT)^[22]^ and developed an integrative analysis platform that enables batch processing of multi-target SMLM data and provides quantitative molecular information. Using this workflow, we investigated the oligomerization of EGFR and the spatiotemporal reorganization of EGFR and Grb2. This imaging and analysis workflow is broadly applicable and can be used to study the organization of other membrane protein assemblies *in situ*.

## Results and Discussion

We first set up a robust imaging workflow for DNA-PAINT super-resolution imaging of EGFR and Grb2 in fixed cells using Exchange-PAINT (**Figure 1A**).^[22]^ To achieve precise axial alignment of the two rounds of image acquisition, we implemented an astigmatic lens into the detection path and used fiducial markers to accurately adjust the imaging plane (**Figure S1, Supporting Note 1**). Using this approach, we visualized the nanoscale distribution of EGFR and Grb2 in untreated HeLa cells and in cells treated with EGF for 1, 5, and 15 min (**Figure 1B**). Protein clusters were isolated from nonspecific signals using density-based spatial clustering of applications with noise (DBSCAN) in combination with a set of kinetic filters (see Methods). The experimental localization precision was calculated to be 10.9 ± 2.3 nm (EGFR) and 12.4 ± 2.4 nm (Grb2), employing a nearest neighbor-based analysis (NeNA).^[23]^

**Figure 1:**
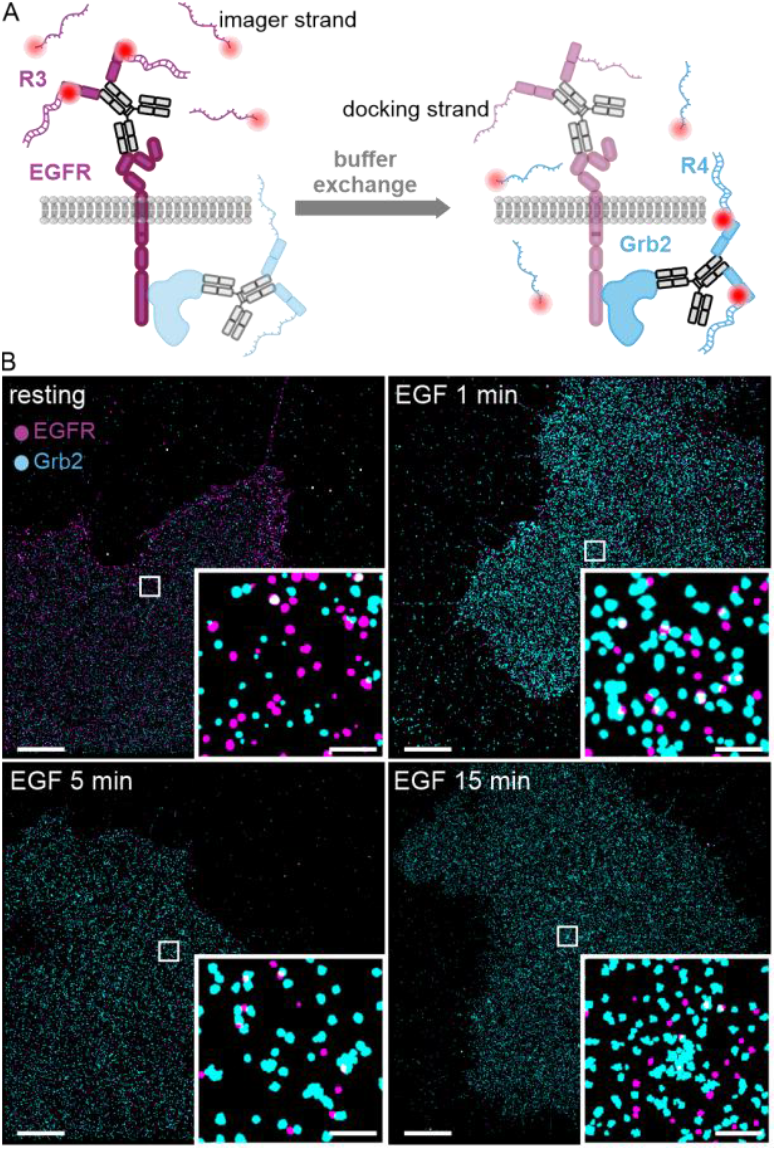
Multiplexed DNA-PAINT super-resolution imaging of EGFR and Grb2 in resting HeLa cells and at different time points of EGF stimulation. A) Scheme of the Exchange-PAINT imaging workflow. EGFR and Grb2 were labeled with preincubated primary antibodies and secondary nanobodies coupled to R3 and R4 DNA docking strands,^[24]^ respectively. After imaging the first target (EGFR), the R3 imager strand was removed by washing, and the second imager strand R4 was added for imaging Grb2. B) DNA-PAINT images of EGFR and Grb2 in the absence of a ligand (resting state) and following EGF stimulation for 1, 5, and 15 min. EGFR is shown in magenta, Grb2 in cyan. Scale bars 5 µm, zoom-ins 0.5 µm.

We next analyzed the density of protein clusters at the plasma membrane of HeLa cells. We observed a significant decrease in the density of EGFR clusters from 7.8 ±2.4 clusters/µm^2^ in the resting state to 4.3 ±1.5 clusters/µm^2^ after 1 min of EGF treatment, to 3.4 ±0.8 clusters/µm^2^ after 5 min stimulation, and to 2.6 ±0.6 clusters/µm^2^ after 15 min stimulation (**Figure 2A**). The observed decrease in EGFR at the plasma membrane upon EGF stimulation is consistent with previous studies.^[25–27]^ The Grb2 cluster density at the plasma membrane in the absence and presence of EGF did not change significantly (**Figure 2A**). This rather constant cluster density of Grb2 is in line with a previous report^[28]^ and can be attributed to continuous and dynamic recruitment and internalization of Grb2.

**Figure 2:**
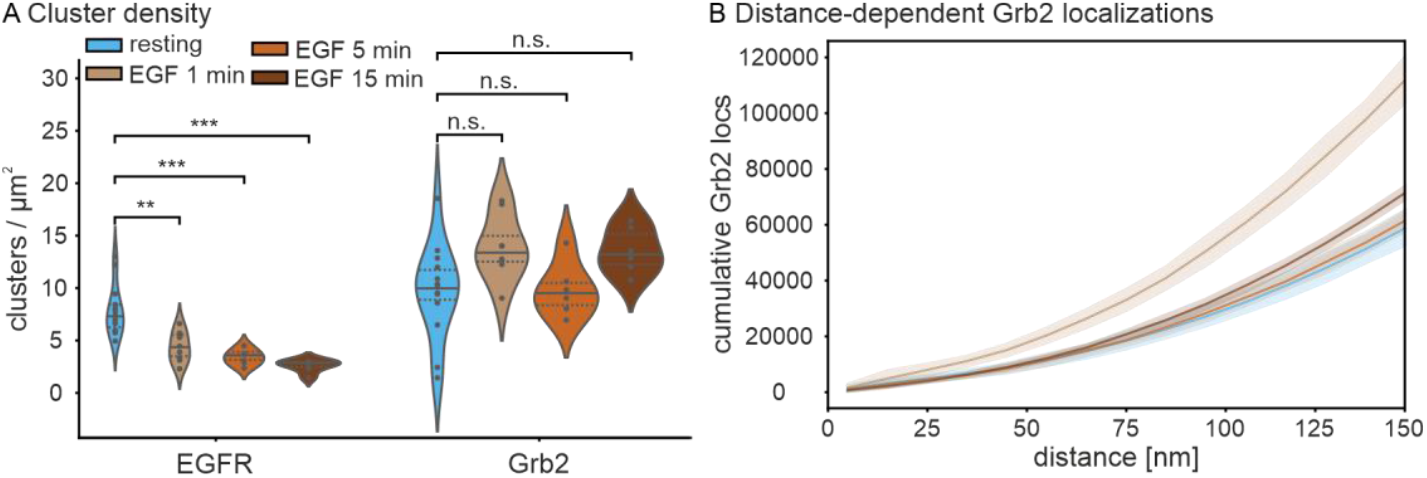
Cluster analysis of EGFR and Grb2 in resting and EGF-stimulated HeLa cells treated for 1, 5, and 15 min. A) Protein cluster density distribution. Points denote densities determined per cell, solid lines indicate the median, and dotted lines mark the 1st and the 3rd quartiles. Statistical significance: ns, not significant (*p >* 5.0 × 10^-2^); ** (1.0 × 10^-3^ < *p* ≤ 1.0 × 10^-2^; *** (1.0 × 10^-4^ < *p* ≤ 1.0 × 10^-3^). B) Cumulative distribution of Grb2 localizations within 150 nm from EGFR cluster centers. Bin size is 10 nm. The shaded band indicates the standard deviation (± SD) of the cumulative curves per bin across individual images. N = 14 (resting), 8 (1 min EGF), 6 (5 min EGF), and 6 cells (15 min EGF).

Next, we investigated whether Grb2 undergoes spatial reorganization relative to EGFR following EGF stimulation. We quantified Grb2 recruitment to EGFR by comparing the cumulative Grb2 localizations within a distance range of 0 to 150 nm from EGFR (**Figure 2B**). The localizations of Grb2 upon 1 min EGF stimulation increased predominantly in comparison to other EGF treatment times and the resting state (p-value = 0.029). In addition, the Grb2 localizations exhibited a modest increase following 5 and 15 min of stimulation compared to the resting state. Our observation is in line with other *in vitro* and *in situ* studies,^[27,29–31]^ and provides evidence of the direct interaction between EGFR and Grb2 at the plasma membrane at the single-molecule level. The number of Grb2 localizations near EGFR decreased again after 5 and 15 min EGF stimulation, possibly due to the accumulation of Grb2 and EGFR in endosomes, which are outside of our observation volume.^[30]^

EGF stimulation induces EGFR dimerization.^[5]^ To assess this in a time-dependent manner, we performed quantitative DNA-PAINT (qPAINT)^[32]^ to determine the oligomerization state of EGFR at different time points of EGF treatment. qPAINT extracts molecule numbers from the binding kinetics between imager and docking strands (**Figure 3A**). The mean dark time or τ_d_ is inversely proportional to the influx rate ξ of imager strands, which is determined by the association rate constant k_on_ and the imager strand concentration C_i_ (ξ = k_on_ × C_i_). Therefore, 1/τ_d_ is linearly proportional to the number of available binding sites under identical imaging conditions (N = 1/(τ_d_ × ξ)).^[32,33]^ Thus, clusters with more binding sites exhibit larger 1/τ_d_ values, enabling estimation of the relative oligomerization state.

**Figure 3:**
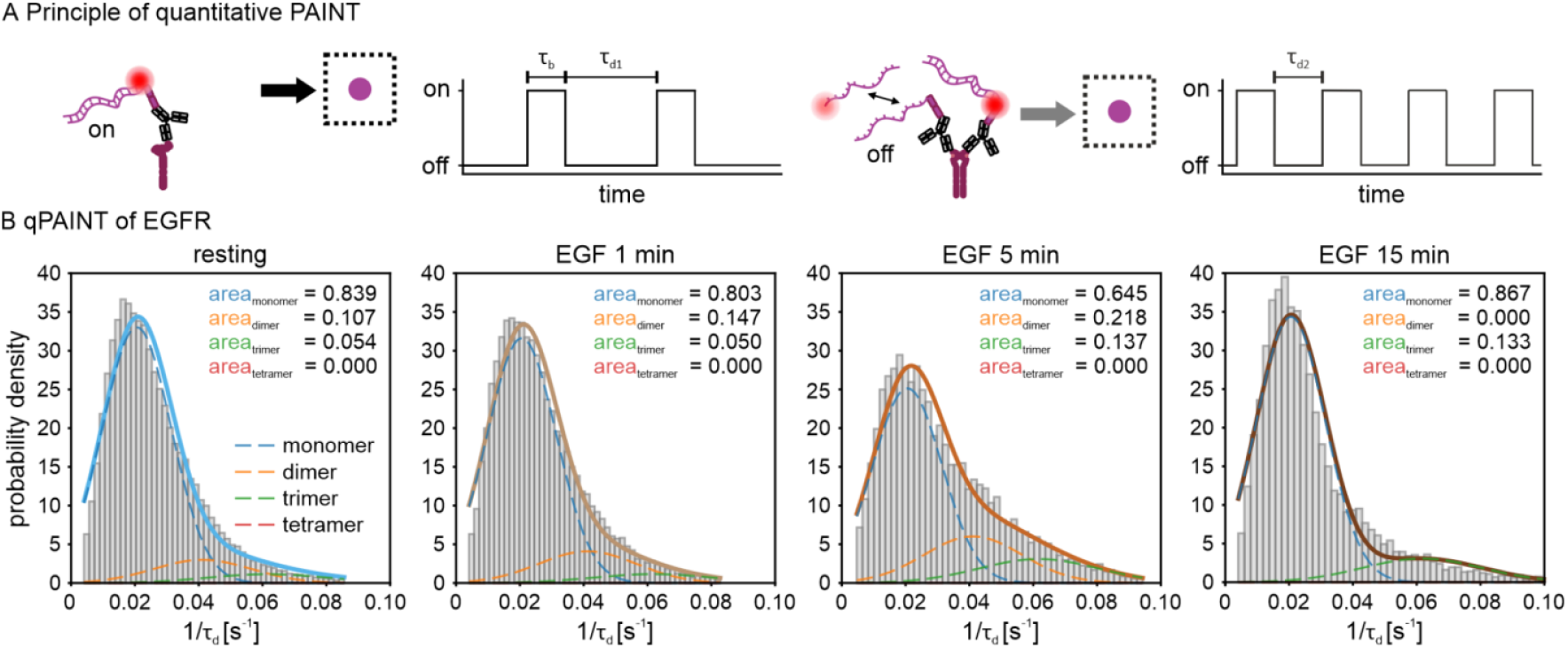
Quantitative analysis of molecule numbers in EGFR clusters under resting and EGF-stimulated conditions. A) Schematic representation of the qPAINT principle. Transient binding between imager and docking strands gives rise to alternating dark periods (τ_d_, when the imager strand is not bound) and bright times (τ_b_, when the imager strand is bound). Because τ_d_ is inversely proportional to the number of available docking strands, clusters with a higher oligomeric state exhibit shorter τ_d_ values due to more frequent binding events. B) Frequency distributions of 1/τ_d_ values derived from individual intensity traces for resting and EGF-stimulated EGFR. Histograms are shown on a probability density scale where the area of all bars sums to 1. The distributions were fitted with a sum of four Gaussian functions (solid lines) corresponding to monomeric, dimeric, trimeric, and tetrameric receptor populations (dashed lines). The area under each curve reflects the estimated fraction of molecules in the respective oligomeric state. N = 14 (resting), 8 (1 min EGF), 6 (5 min EGF), and 6 cells (15 min EGF).

Approximation of the 1/τ_d_ distributions with a sum of Gaussian functions revealed the relative fractions of clusters containing monomeric or higher-order oligomeric EGFR (**Figure 3B**) (Methods). While resting cells showed mainly one population, which we attribute to monomeric EGFR, the fraction of monomeric EGFR decreased slightly after 1 min of EGF stimulation and reached its minimum after 5 min. Cells that were treated for 15 min with EGF showed monomer levels similar to values observed in resting cells. Conversely, EGF stimulation led to an increase in both dimeric and higher-order oligomeric EGFR fractions. Interestingly, after stimulating the receptor for 5 min with EGF, a discernible population of higher oligomeric states began to appear, which lends support to the previously described tetrameric EGFR structure.^[15,34,35]^ For Grb2, we observed that there was a steady accumulation of proteins within individual Grb2 clusters, and this persisted throughout 15 min of EGF stimulation (**Figure S2**).

We next sought to establish an orthogonal and alternative analysis of multi-target DNA-PAINT data that can reveal the different activation states of cells. Uniform Manifold Approximation and Projection (UMAP)^[36]^ has become a widely used dimensionality reduction approach for visualizing high-dimensional biological data and revealing biologically meaningful information.^[37–40]^ We selected UMAP over t-SNE^[41]^ or principal component analysis (PCA)^[42]^ to study the spatiotemporal reorganization of EGFR and Grb2 upon ligand stimulation since it preserves both local similarities between cells with comparable nanoscale features and global relationships between different conditions.^[36]^ To visualize condition-dependent differences in EGFR and Grb2 organization, we projected 44 features (see Methods) describing protein properties into a 2D UMAP representation. The embedded data for EGFR and Grb2 were averaged per cell and color-coded per condition (**Figure 4A, B**). *k*-means clustering revealed four clusters, each containing a mixture of resting and EGF-stimulated conditions but exhibiting distinct condition-dependent enrichment (**Figure 4C, D**). It is noteworthy that cluster 1 is predominantly composed of the resting condition. Cluster 2 comprises a heterogeneous population of cells, encompassing both resting and stimulated (EGF 1 and 5 min) cells. Clusters 3 and 4 exclusively contain stimulated cells for EGFR and primarily stimulated cells for Grb2. Interestingly, 1 min EGF-stimulated cells showed the highest heterogeneity and appeared in all clusters as a transitional state, while resting cells and cells stimulated for 15 min with EGF showed the most pronounced differences. Additionally, we noticed a clear separation between EGFR and Grb2 clusters reflecting their distinct functional roles (**Figure S3**). Overall, the clustered data revealed condition-dependent trends, emphasizing nanoscale reorganization of the receptor upon EGF activation.

**Figure 4:**
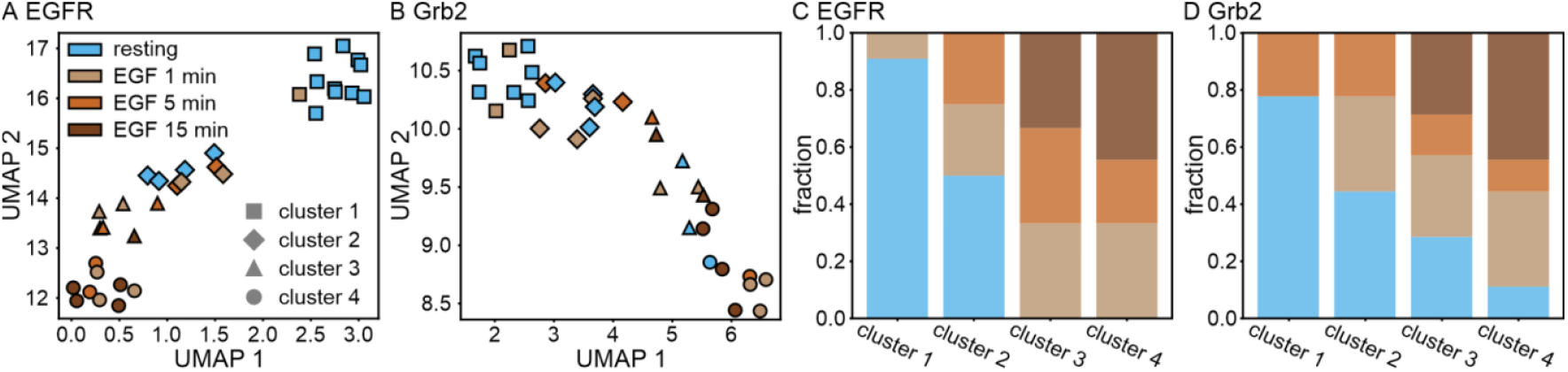
UMAP projection of A) EGFR and B) Grb2 features across resting and EGF-stimulated cells. A total of 44 features were included. Each point represents a cell. Points are color-coded based on condition and grouped via k-means clustering. N = 14 (resting), 8 (1 min EGF), 6 (5 min EGF), and 6 cells (15 min EGF). The barplots represent the distribution of conditions within each cluster of C) EGFR and D) Grb2, respectively.

## Conclusion

We report the visualization and quantification of the nanoscale organization of the transmembrane receptor EGFR and its adaptor protein Grb2 in HeLa cells using quantitative DNA-PAINT at the single-molecule level. This analysis was performed *in situ* under resting conditions and following EGF stimulation at multiple time points. We observed a stimulation-dependent decrease in EGFR density at the plasma membrane, while Grb2 density remained largely constant but underwent pronounced spatial reorganization relative to EGFR. Furthermore, analysis of oligomeric states within individual clusters revealed a ligand-induced shift toward higher-order oligomers of both EGFR and Grb2. Spatiotemporal differences between stimulation conditions were further corroborated by UMAP embedding and k-means clustering of multidimensional single-molecule features. In summary, we present a versatile single-molecule data analysis workflow that is readily transferable to other receptor systems, enabling spatiotemporal characterization of membrane protein assemblies *in situ*.

## Experimental Section

### Cell culture

The human cervical cancer cell line HeLa (# ACC 57, DSMZ, Braunschweig, Germany) was cultured in growth medium consisting of Dulbecco’s modified Eagle’s medium (DMEM) (# 11574486, Gibco, Life Technologies, Waltham, MA, USA) supplemented with 10% fetal bovine serum (# 35-079-CV, Corning Inc., Corning, NY, USA), 1 unit/mL penicillin and 1 µg/mL streptomycin (Gibco, Life Technologies), and 1% v/v GlutaMAX (# 35050-038, Gibco, Life Technologies). The cells were incubated at 37 °C with 5% CO_2_ and were passaged every 3–4 days. HeLa cells were seeded in ibidi µ-Slide (# 80607, ibidi GmbH, Gräfelfing, Germany) coated with PLL-PEG-RGD (poly-l-lysine-grafted polyethylene glycol modified with a CGRGDS peptide)^[43]^ at a density of 1×10^5^ cells/mL for one day of growth.

### Sample preparation

The cells were starved for 2 hours prior to ligand stimulation using growth media without fetal bovine serum. After starvation, EGF (# AF-100-15, PeproTech, Thermo Fisher Scientific, Waltham, MA, USA) was diluted in serum-free medium to an end concentration of 100 ng/mL and then incubated with cells for 1, 5, or 15 min. The cells were fixed using prewarmed 3% formaldehyde (# 28908, Sigma-Aldrich, St. Louis, MO, USA) with 0.25% glutaraldehyde (# G5882, Sigma-Aldrich) in 1x phosphate-buffered saline (PBS, # 14190144, Gibco) and incubated for 15 min at 37 °C. Monoclonal mouse against EGFR (1:50 dilution, # sc-120, Santa Cruz Biotechnology, Texas, USA) and mouse against Grb2 (1:100 dilution, # MA5-34887, Thermo Fisher) were pre-incubated with secondary anti-mouse nanobodies (2-fold molar excess) conjugated to R3 and R4 docking strands (custom-labeled, Massive Photonics, München, Germany), respectively, for 1 h at 4 °C in blocking buffer (1 mM EDTA, 0.02% Tween 20, 0.05% NaN_3_ (all from Sigma-Aldrich), 2% bovine serum albumin (# 9048-46-8, Carl Roth GmbH & Co. KG, Karlsruhe, Germany), 0.05 mg/mL salmon sperm DNA (#AM9680, Invitrogen, Thermo Fisher Scientific) in 1x PBS). The antibody-nanobody complexes were then combined and applied to the cells for 2 h at room temperature. 100 nm gold beads (# A11-100-NPC-DIH-1-100, Nanopartz Inc., Loveland, US) were used as fiducial markers. They were diluted 1:5 in 1x PBS, sonicated for 10 min, and incubated for 15 min, followed by washing with 1x PBS. The cells were post-fixed using 4% formaldehyde in 1x PBS.

### DNA-PAINT imaging

DNA-PAINT imaging was performed on a home-built widefield setup based on a Nikon Eclipse Ti inverted microscope. Excitation was provided by a 561 nm laser (200 mW Sapphire, Coherent Inc., Santa Clara, CA, USA) with laser power modulated via an acousto-optic tunable filter (AOTFnC-400.650-TN, AA Opto Electronic, Orsay, France). To ensure a clean beam profile, the laser was fiber-coupled using a collimator (60FC-4-M6.2-33) into a polarization-maintaining single-mode optical fiber (PMC-E-400RGB), and then re-collimated to a 6 mm full width at half maximum (FWHM) beam (60FC-T-4-M50L-01; all from Schäfter & Kirchhoff GmbH, Hamburg, Germany). The collimated beam was expanded using a telescope (AC255-030-A-ML and AC508-150-A-ML, Thorlabs GmbH, Dachau, Germany) and focused onto the back focal plane of a 100× TIRF oil immersion objective (CFI Apochromat TIRF 100XC Oil, NA 1.49, Nikon, Tokyo, Japan). A motorized mirror (MTS50-Z8, Thorlabs) enabled adjustment of the illumination angle for widefield, HILO, or TIRF imaging modes. Axial focus was stabilized using a Perfect Focus System (Ti-PFS, Nikon), while lateral sample positioning was controlled by a motorized stage (Ti-S-ER, Nikon) in combination with a piezo stage (Nano-Drive, MadCityLabs, USA). Excitation light was introduced into the microscope via a multiband dielectric beamsplitter (zt405/488/561/640rpc, AHF Analysentechnik AG, Tübingen, Germany), which also transmitted emission light into the detection path. Fluorescence was spectrally filtered with a bandpass filter (610/60 ET, Chroma, Bellows Falls, VT, USA) and imaged with an EMCCD camera (iXon Ultra DU-897U-CS0, Andor, Belfast, UK).

Before imaging, the corresponding Cy3B-labeled imager strand (R3 or R4) was diluted to a final concentration of 2 nM in imaging buffer containing 5 mM Tris/HCl (pH 8.0), 75 mM MgCl_2_, and 0.05% Tween-20 (all Sigma-Aldrich), freshly supplemented with an oxygen scavenging and triplet-state quenching system (1× protocatechuic acid (# 37580-25G-F, Sigma-Aldrich), 1× protocatechuate-3,4-dioxygenase from Pseudomonas sp. (# P8279-25UN, Sigma-Aldrich), and 1× (±)-6-hydroxy-2,5,7,8-tetramethylchromane-2-carboxylic acid (Trolox) (# 238813, Sigma-Aldrich)), prepared as described by Schnitzbauer et al.^[44]^ DNA-PAINT imaging was conducted using a 561 nm laser at an excitation intensity of 0.1 kW/cm^2^. All microscope components were controlled via the µManager software.^[45]^ Image acquisition consisted of 20,000 frames per target recorded with the following parameters: 100 ms exposure time, EM gain of 150, 3× preamplifier gain, 10 MHz readout rate, image size of 256×256 pixels, and frame transfer mode activated. Cell positions were saved within µManager to facilitate subsequent multi-target imaging. Bright-field images were acquired before and after each DNA-PAINT acquisition. For Exchange DNA-PAINT experiments, the sample was washed five times with 1× PBS after imaging each target, followed by continuous laser illumination for 5 minutes to ensure complete removal of residual imager strands. Afterwards, the second imager strand was added. In total, at least three independent samples for each condition were prepared and acquired as described above.

### Data analysis

Image processing was performed using Picasso software (version 0.7.3).^[44]^ First, single emitters in each frame were localized by fitting the Linear-Quadratic (LQ) Gaussian fit with a min. net gradient of 20,000, a box side length of 7, an EM gain of 150, a baseline of 88.3, a sensitivity of 0.95, and a pixel size of 158 nm. Next, drift correction was performed using fiducial markers. Localized single-molecule events were filtered for the width of the point spread function (PSF). Localizations appearing in multiple consecutive frames were merged using parameters derived from nearest neighbor-based analysis,^[23]^ with a merging radius set to four times the NeNA value and a maximum dark time of one frame. Next, DBSCAN clustering was performed using 1x NeNA and a minimum of 7 min localizations. The identified clusters were further filtered based on the mean frame time within a range of µ-2*δ*to µ+2*δ*, where µ represents the average mean frame time and *δ*is the standard deviation (1,500-8,000 frames).

Clustered localization data were processed using a custom-written Python-based analysis pipeline capable of handling cluster-center information from multiple .*hdf5* files across an arbitrary number of proteins and experimental conditions. The analyzed cell area was defined using pre-generated masks, and cluster and localization densities were calculated per µm^2^. Nearest-neighbor relationships and corresponding distances were computed for self- and, optionally, cross-protein interactions. To compare the cumulative distributions of Grb2 localizations relative to EGFR, the area under the curves was calculated, and each condition was compared to the resting state.

qPAINT analysis was performed with optional calibration using a reference signal, such as antibodies nonspecifically bound to the glass surface adjacent to the imaged cell. For qPAINT, the distribution of inverse dark times (1/τ_d_) was fitted using a Gaussian model, in which the peak position corresponding to the monomer population was fixed based on the resting condition, where proteins are expected to predominantly exist as monomers.

For dimensionality reduction, 44 features capturing nanoscale organization and protein behavior were extracted from each protein and projected into a two-dimensional space using UMAP (version 0.5.5).^[36]^ These features comprised cluster-level properties, localization-level densities, binding kinetics parameters derived from qPAINT analysis, and spatial neighborhood organization. For each feature, summary statistics (mean, standard deviation, minimum, and maximum) were computed per cell to capture both central tendencies and heterogeneity. The resulting coordinates were clustered using the k-means algorithm^[46]^ from the scikit-learn library (version 1.3.0) to identify distinct sub-groups. The number of clusters k was set equal to the number of experimental conditions.

Statistical tests were performed to examine significant differences between conditions. The statistical analysis was conducted using the non-parametric Mann-Whitney U test from the *statannot* library (version 0.2.3).

## Supporting Information

The authors have cited additional references within the Supporting Information.^[47]^

## Acknowledgement

This work was supported by the Deutsche Forschungsgemeinschaft (DFG), grants CRC 1507 and GRK 2566 (iMOL). We are grateful to Petra Freund for support with cell culture, and to the members of the SMB team for helpful discussions and feedback.

