## Supplemental Information for "Quantitative mapping of nanoscale EGFR–Grb2 assemblies by DNA-PAINT"

### Supporting Note

#### Supporting note 1

Exchange DNA-PAINT is a powerful technique for visualizing multi-protein assemblies in cells.[1] In this approach, different targets are labeled with sequence-orthogonal docking strands and detected in sequential imaging rounds, interspersed with washing and exchange of the corresponding imager strands. Exchange DNA-PAINT can be performed using the same fluorophore, thereby avoiding crosstalk between detection channels and chromatic aberrations. However, this multiplexing strategy requires precise alignment of the lateral (x–y) position and the axial (z) focal plane to be identical across all acquisition rounds. While this is readily achieved for *in vitro* samples, such as DNA origami structures or proteins immobilized on coverslips, it becomes more challenging when applied to cellular systems.

When studying the spatial relationship between transmembrane proteins and cytoplasmic effector molecules in cells, the intrinsic curvature of the plasma membrane[2] makes accurate axial alignment essential (Figure S1). Washing steps between consecutive rounds may result in axial (z-direction) displacements of several hundred nanometers, leading to an artificially reduced apparent colocalization rate (Figure S1B). Therefore, in addition to lateral alignment, we implemented astigmatism-based axial alignment before each data acquisition round to correct for z-direction offsets. We achieved this by using gold beads as fiducial markers and by inserting an astigmatism lens during z-plane adjustment between imaging rounds (Figure S1C-E). This axial alignment procedure resulted in a pronounced increase in the measured colocalization rate (Figure S1A).

### Supporting Figures

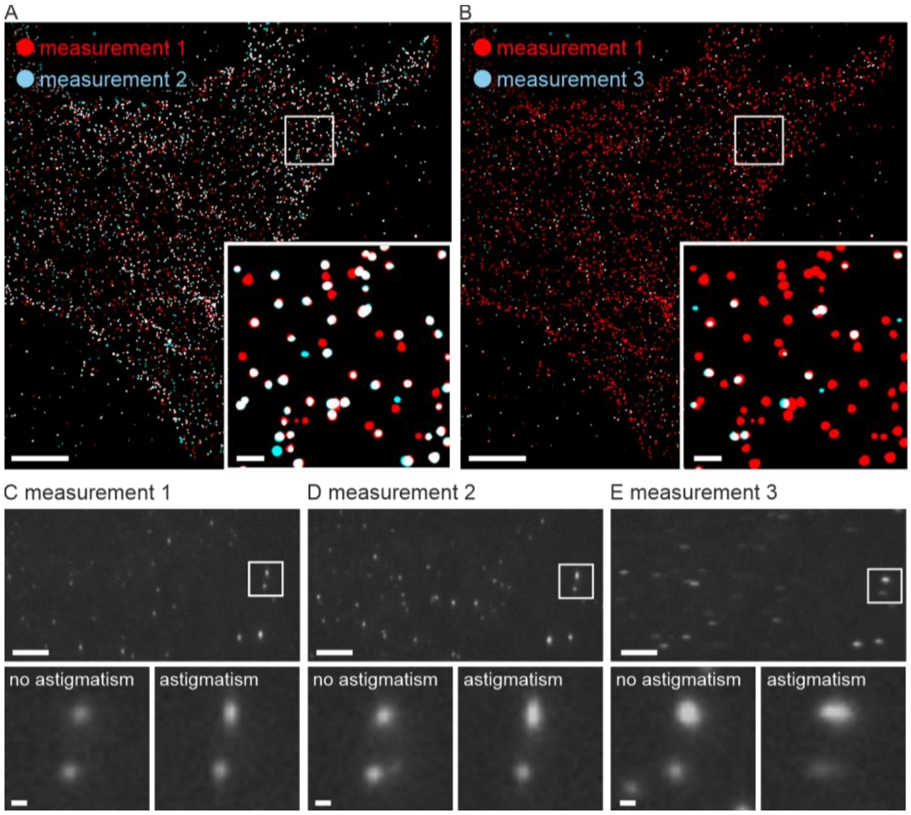

**Figure S1**: Axial alignment in multiplexed DNA-PAINT with and without astigmatism. A) Astigmatism-based axial alignment results in high colocalization (~84%) of EGFR localizations obtained from consecutive acquisition rounds. B) In the absence of astigmatism-based axial alignment, a substantially reduced colocalization to 15% of EGFR localizations is observed between consecutive acquisitions. C–E) Upper panels show overview images of fiducial markers acquired during measurement rounds 1–3 with an astigmatic lens inserted into the detection path. Lower panels show zoomed-in views of fiducial beads with or without astigmatism-based axial alignment. Scale bars 5 μm, zoom-ins 500 nm.

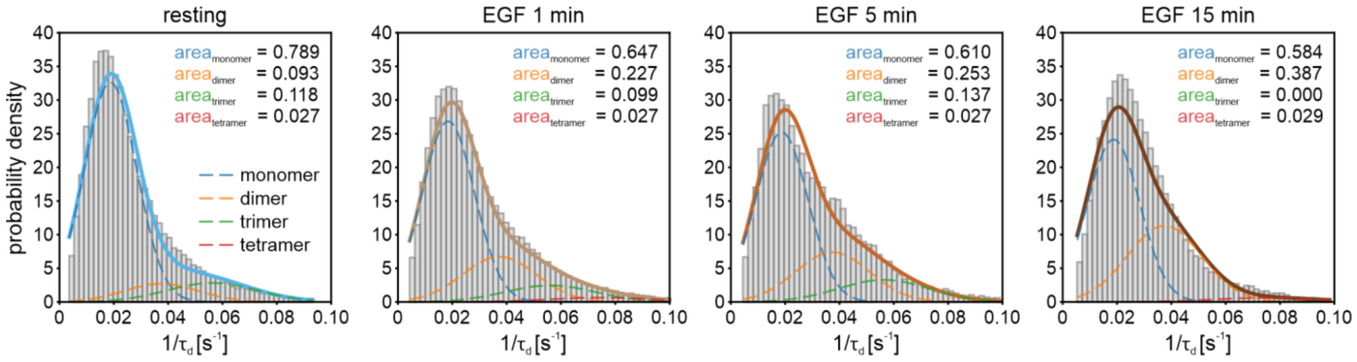

**Figure S2**: qPAINT analysis of Grb2 clusters under resting and EGF-stimulated conditions. Frequency distributions of $1/\tau_d$ values derived from individual intensity traces of Grb2 for resting and EGF-stimulated cells. The distributions were fitted with a sum of four Gaussian functions (solid lines) corresponding to monomeric, dimeric, trimeric, and tetrameric protein populations (dashed lines). The area under each curve reflects the estimated fraction of molecules in the respective oligomeric state.

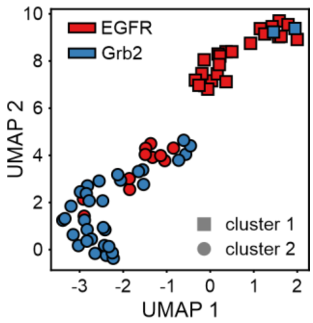

**Figure S3**: UMAP projection of EGFR and Grb2 features. The resting and stimulated data were pooled for EGFR (red) and Grb2 (blue), respectively. A total of 44 features were included. Each point represents a cell. Data points were grouped via k-means clustering. N = 34 cells.
